# MBIOME: A comprehensive, reproducible, and open-source workflow for amplicon-based microbiome data analysis

**DOI:** 10.64898/2026.06.25.734448

**Authors:** Miriam Gorostidi-Aicua, Ane Otaegui-Chivite, Aitor Zabala, Laura Moles, David Otaegui

## Abstract

Microbiome analysis has become a pivotal tool in understanding the role of microbiota in human health and disease. However, the lack of standardized workflows, together with the limitations of proprietary software solutions, hampers reproducibility and flexibility. Here, we present *Mbiome*, an open-source, user-friendly and automated workflow designed to streamline amplicon-based microbiome analysis. Built upon QIIME2, *Mbiome* supports both bacterial (16S rRNA) and fungal (ITS) profiling, and is compatible with raw fastq files generated by Ion Torrent (IT) and Illumina (IL) sequencing platforms.

The workflow guides users through an interactive setup process via a simple configuration file, enabling researchers with minimal bioinformatics experience to perform comprehensive analyses without writing code. Once configured, *Mbiome* automates major steps including quality control, taxonomic assignment, *α* and *β*-diversity analyses, functional predictions (via q2-metnet), and customizable visualizations and statistical analyses.

*Mbiome* has been validated using real-world datasets from multiple sclerosis research projects, performing a comparison between different microbiome analysis approaches, including 16S hypervariable region reconstruction, amplicon-based strategies, and cross-platform sequencing (IT and IL), as well as against results obtained with Ion Reporter (IR) commercial software. This evaluation demonstrated its versatility and effectiveness across different sequencing platforms. Moreover, *Mbiome* provided improved flexibility, transparency, and taxonomic resolution compared to IR. By combining accessibility, reproducibility, and cross-platform compatibility, *Mbiome* lowers the barrier to microbiome data analysis and facilitates high-quality, standardized workflows in both research and applied settings. *Mbiome* is publicly available at https://github.com/MGorostidi/mbiome.

## Introduction

Microbiome research has emerged in the last decade, uncovering key interactions between microbial communities and host physiology (Lloyd-Price et al., 2016; iHMP, 2014). Several studies have demonstrated the effect of microbiota on health and disease, especially regarding gut microbiota, which constitutes the most complex and extensive microbial ecosystem in the human body (Moles and Otaegui, 2020; Chen et al., 2021; Afzaal et al., 2022)

In this context, high-throughput sequencing technologies have become indispensable for investigating these complex microbial ecosystems. Advances in sequencing have opened the door to multiple strategies to study microbiota, from whole-metagenome shotgun sequencing to targeted approaches that focus on specific phylogenetic markers (Knight et al., 2018; Caporaso et al., 2010).

Among the various sequencing approaches, targeted amplicon sequencing—especially focused on the 16S rRNA gene for bacteria and archaea, and ITS intergenic regions for fungi—remains the method of choice for taxonomic profiling of microbial environments. The targeted approach provides multiple advantages, such as reduced sequencing costs by focusing only on specific regions, enabling robust multiplexing of hundreds of samples in a single run (quicker analysis), higher sequencing depth per target region resulting in improved taxonomic resolution, and low-biomass requirement (Liu et al., 2020; Yang et al., 2016).

Regarding amplicon sequencing workflow, this comprises the following key stages (Liu et al., 2020): sample collection and DNA extraction, amplification of specific phylogenetic marker regions using PCR, preparation of sequencing libraries and sequencing by high-throughput techniques (typically on platforms like Ion Torrent (IT) or Illumina (IL)). Here, the bioinformatic processing starts: the resulting raw reads undergo quality control and filtering to remove low-quality sequences and artifacts; then, reads are either denoised to resolve exact amplicon sequence variants (ASVs) or clustered into operational taxonomic units (OTUs), followed by chimera removal to eliminate sequencing artifacts. Representative sequences are subsequently assigned taxonomic classifications using curated reference databases. Finally, community structure is explored using diversity analyses (*α* and *β*-diversity), differential abundance testing, and visualization techniques to interpret ecological patterns within the microbiota.

While laboratory and sequencing protocols have become increasingly streamlined, a persistent challenge remains: the lack of standardized, transparent, and flexible computational pipelines for microbiome analysis. This situation leaves researchers uncertain about where to start, which parameters or algorithms to use, and makes it difficult to reproduce and compare results between studies. Furthermore, previous studies, have already addressed dramatic differences in the resulting microbial community composition depending on the pipeline followed (Straub et al., 2020). Moreover, the information available related to the analysis usually depends on the sequencing platform used, habitually IL, leaving almost out of scope the analysis of samples sequenced with other equipment, such as IT.

Thermo Fisher Scientific provided a commercial software called Ion Reporter (IR) for microbiome analysis of IT sequencing files. These can be directly chosen from the sequencing equipment to be analyzed with the software. The software offers a user-friendly interface, however, it functions as a “black box”, limiting parameter customization. The dependence on external technical support, and lack of analytical transparency motivated the development of an in-house solution.

Open-source platforms like QIIME2 (Bolyen et al., 2019) have emerged as the gold standard in microbiome analysis due to their modularity and strong community support. Nonetheless, QIIME2’s command-line interface poses a barrier for researchers without bioinformatics expertise, especially when dealing with complex datasets such as those generated by the Ion 16S Metagenomics Kit, which pools multiple hypervariable regions in single fastq files.

To address some of these challenges, we developed Mbiome, a comprehensive and modular bioinformatics workflow for amplicon sequencing analysis built upon QIIME2. *Mbiome* covers all major steps from initial fastq files through to publication-ready figures and results tables. It is designed to maximize reproducibility, transparency, and ease of use, integrating state-of-the-art quality control, denoising, taxonomic classification, diversity estimation, functional predictions, and visualization modules. Around 250 stool samples were used from current projects in our lab to demonstrate *Mbiome*’s utility in real life (Gorostidi-Aicua et al., 2024; Moles et al., 2024; Otaegui-Chivite et al., 2025). In addition, ten stool samples were used to compare the four microbiome analysis pipelines implemented in *Mbiome* and to benchmark its performance against an established workflow, the commercially available software IR. By making the workflow and all scripts fully available, we aim to provide the scientific community with a robust tool for standardized, reproducible amplicon sequencing analysis, democratizing access to advanced bioinformatics workflows for the broader research community.

## Results

### Overview of Mbiome

The primary outcome of this project is the development of *Mbiome*, an internally designed bioinformatics workflow for amplicon sequencing analysis. This tool has been progressively refined over the past years through extensive development efforts.

#### Workflow Components and Analytical Pipelines

*Mbiome* is a comprehensive and automated workflow built upon QIIME2. It covers all major steps of microbiome analysis, from raw fastq files to publication-ready figures and result tables, and is designed to be accessible even to researchers without prior bioinformatics expertise. The overall structure of *Mbiome* (v1.0.0) is illustrated in Figures 1 and 2.

**Figure 1.**
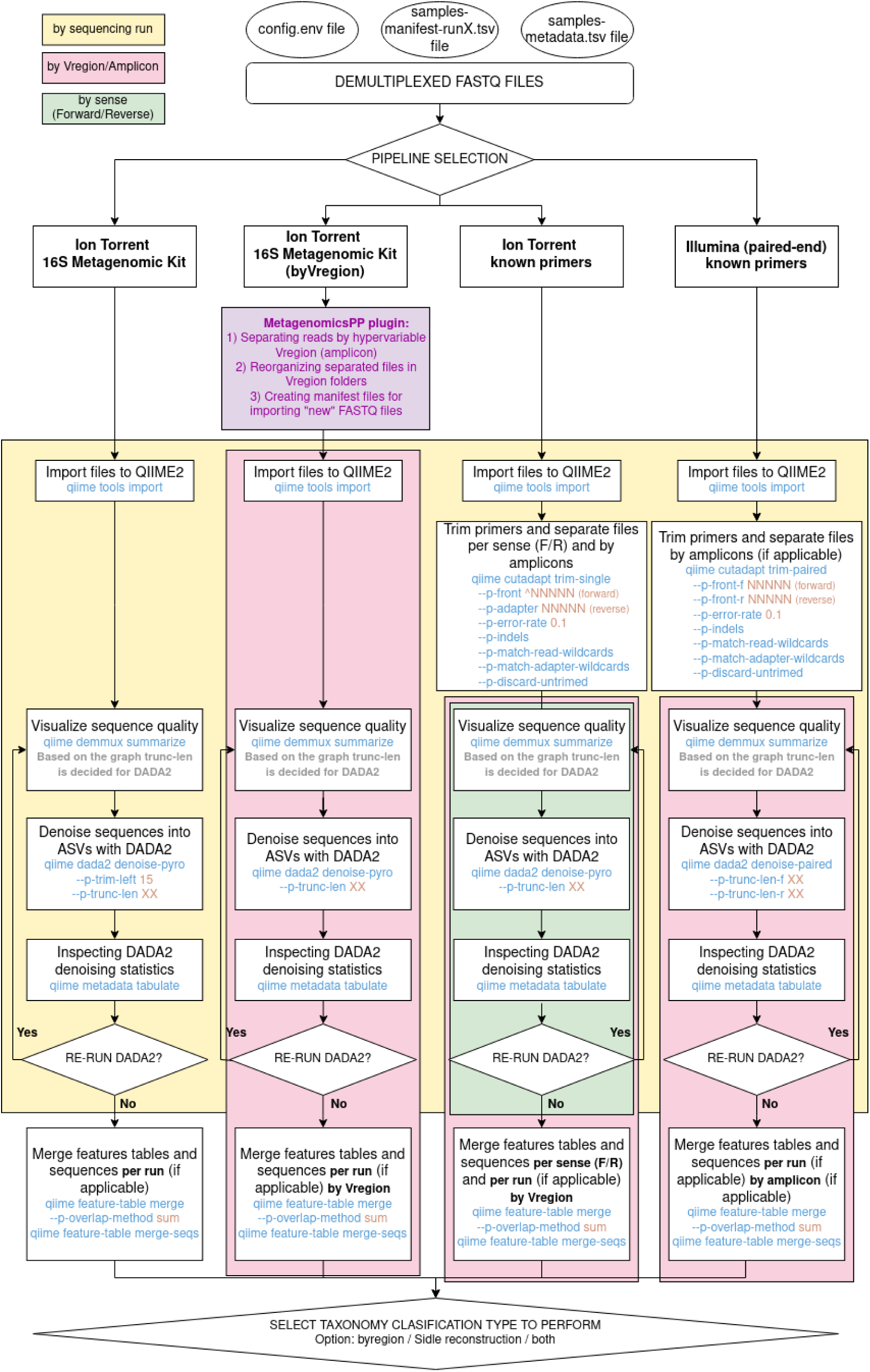
Workflow of the *Mbiome* pipeline.

**Figure 2.**
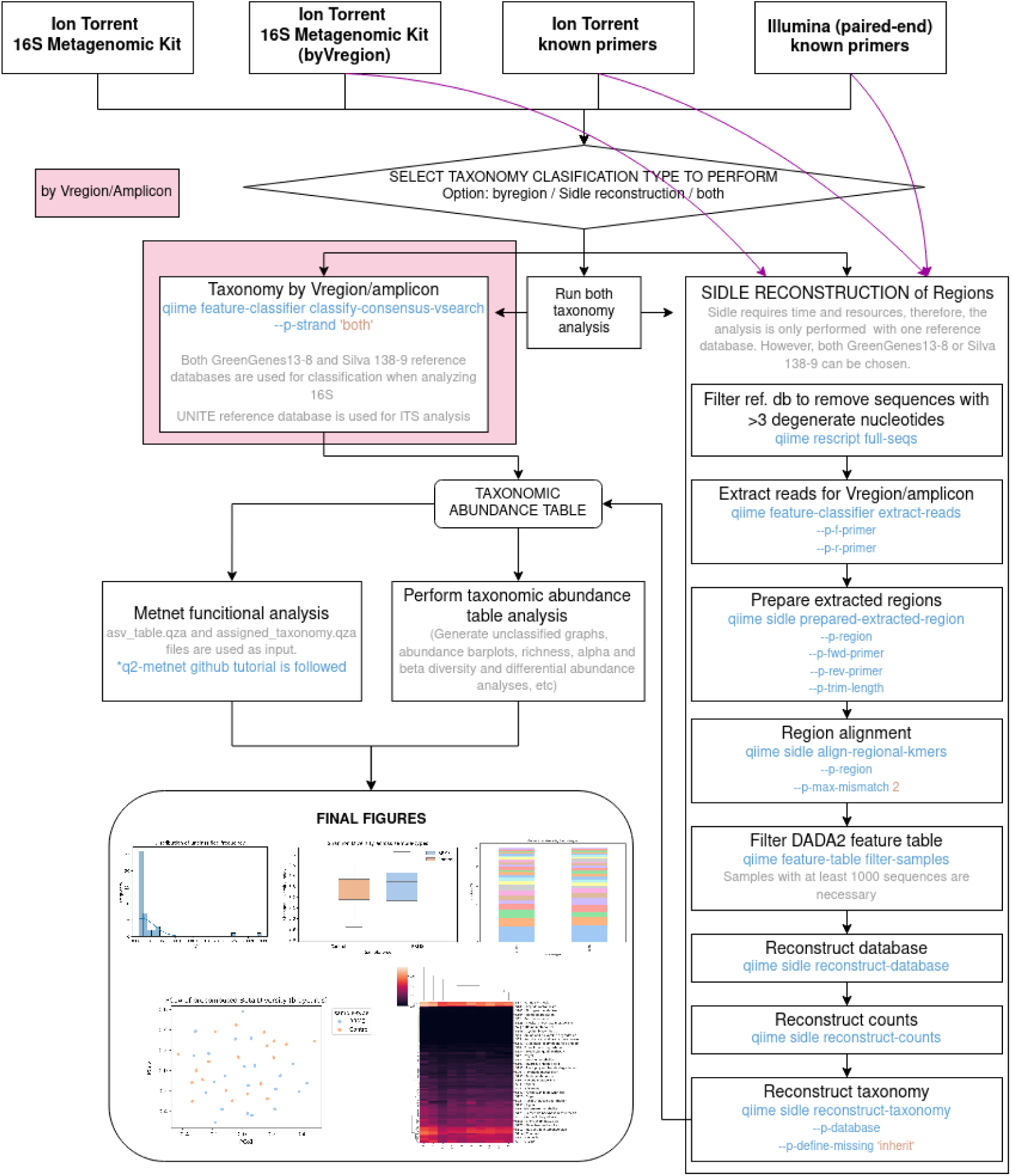
Workflow of the *Mbiome* pipeline (second part).

As shown in Figure 1, *Mbiome* integrates multiple internal pipelines. This modular design ensures that each experiment is processed with parameters tailored to its specific characteristics, including:

- Sequencing platform (e.g., IT or IL)
- Amplification strategy (commercial kit or custom primers)
- Sequencing run configuration (single or multiple runs)
- Targeted analysis type (16S rRNA or ITS)

To facilitate and automate the analysis, users fill a configuration file that contains all the relevant information about the experiment. This file enables *Mbiome* to execute the appropriate workflow based on the specified parameters. Table 1 summarizes the parameters included in this configuration file, which is unique for each experiment.

**Table 1.** Glossary of the parameters recollected for the configuration file.

| Parameters | Description |
| --- | --- |
| EXPERIMENT_NAME | Name of the experiment folder created in ./mbiome/EXPERIMENTS folder. |
| SEQUENCER | Sequencing platform used in the experiment (Options: IT (Ion Torrent) / IL (Illumina)). |
| SEQUENCING_RUNS | Number of sequencing runs that the experiment includes (integer). |
| AMPLIFICATION | Amplification type performed in the experiment (Options: Primers / Kit). If the primers sequences are known, "Primers" is selected, if the Ion 16S Metagenomic Kit was used, instead, "Kit" option is chosen. |
| SPLIT_BY_REGION | This parameter is for the experiments that used 16S Metagenomic Kit, which includes different regions, for deciding if a splitting of regions wants to be performed. If splitting is not desired or primers were used for amplification, 'No' should be selected. (Options: Yes / No). |
| AMPLICONS | Name of the amplicons amplified (any name can be given to the amplicon, for example: "amp1" / "V3V4" / "V2,V3,V67"). If the Ion 16S Metagenomic Kit was used this parameter will be empty, since all the Vregions will be analyzed. |
| _fwd_primer | Here the Forward primer sequences need to be defined, one per amplicon used. If AMPLICONS="V2,V3,V8", then V2_fwd_primer, V3_fwd_primer and V8_fwd_primer need to be included in config file. |
| _rev_primer | Here the Reverse primer sequences need to be defined, one per amplicon used. If AMPLICONS="V2,V3,V8", then V2_rev_primer, V3_rev_primer and V8_rev_primer need to be included in config file. |
| TAXA_ANALYSIS | Type of taxonomic characterization to perform: by region, SIDLE reconstruction (described in Methods section) or both (Options: byregion, sidle or both). |
| ANALYSIS_TYPE | Analysis type, between bacterial or fungal kingdoms (Options: 16S / ITS). |
| SIDLE_REF_DB | SIDLE reconstruction requires a considerable amount of time and resources, so only one database will be selected for performing this step (Options: Silva / GG (GreenGenes)). |
| AMPLICONS_ANALYSIS | Once the abundance table is obtained, the analysis of this can be performed. It is possible to analyze specific amplicons or regions individually or collectively by specifying the desired amplicon(s). For example, if the workflow was executed with AMPLICONS="V2,V3,V8" but only the abundance analysis of a single region is required, the parameter can be set to the target region, such as AMPLICONS_ANALYSIS="V2". |
| TAX_LEVEL_ANALYSIS | Taxonomic level in which the analysis needs to be performed (Options go from Kingdom to Genus: Kingdom, Phylum, Class, Order, Family, Genus). |
| GROUP_METADATA_COLUMN | Metadata group column to analyze in the taxonomic abundance analysis (The name entered must match exactly the corresponding column name in the metadata file). |
| FUNC_ANALYSIS | If a functional analysis using q2-metnet wants to be performed, this parameter need to be filled with a "Yes" (Options: "" / Yes). |
| METNET_WORKFLOW_TYPE | Choose the abundance table in which Metnet funtional analysis needs to be performed (Options: Region / Sidle) (For IT-kit pipelines this parameter won't be necessary). |
| METNET_WORKFLOW_REGIONS | If the workflow type chosen is "Region", but the functional analysis is only necessary for selected regions, specify them (Example: "V2,V3V4,V6") |
| METNET_TAX_LEVEL | Select the Taxonomy level to perform the analysis on (Options: k / p / c / o / f / g) |
| METNET_GROUP | Metadata column that contain the 2 groups for comparative analysis (Example: "sex"). |
| METNET_CONDITION_GROUP | Name of the group that refers to "condition" or disease group. Or, failing that, the name of one of groups. |
| METNET_CONTROL_GROUP | Name of the group that refers to "control". Or, failing that, the name of the other group. |

Once the configuration file, the QIIME2 manifest.tsv (for file import), and the metadata.tsv files (sample-related information) are created (described in Methods section), the analysis can be initiated. The workflow selected for each experiment depends on its specific characteristics. As depicted in Figure 1, *Mbiome* comprises four pipelines that share the same overall structure, but with subtle yet critical differences:

1. **IT Metagenomics Kit:** This pipeline processes data generated using the 16S Metagenomic Kit for DNA amplification and the IT sequencing platform. Because the proprietary primer sequences of this Kit are unknown, primer trimming (15 bp) is performed during the DADA2 step. After quality control and merging of feature tables across runs (if applicable), taxonomic classification is conducted. As no splitting by 16S V hypervariable region (hypervariable regions are described in Figure 14) is possible, only one abundance table is produced, and the taxonomic analysis follows a region-based approach (no SIDLE reconstruction is available—described in Methods section).
2. **IT Metagenomics Kit (by V region):** This pipeline also processes IT 16S Metagenomic Kit data, but applies the MetagenomicsPP plugin to remove proprietary primer sequences and reorganize reads by V region (Maki et al., 2023). Subsequent importing, DADA2 processing, and merging are performed separately for each region (shown in red color in the workflow diagram). Two taxonomic analysis strategies are available: (i) independent region-based analysis or (ii) global taxonomic profiling through SIDLE reconstruction.
3. **IT known primers:** In this pipeline, IT sequencing is used with known commercial primers. Primer removal and splitting by V region is performed with cutadapt following recommended parameters (Maki et al., 2023). Since forward and reverse primers generate distinct amplicons, cutadapt and DADA2 are executed separately for each orientation (shown in green color in the workflow diagram). Depending on whether a single or multiple amplicons were amplified, the workflow performs either a direct analysis or a split by region/amplicon. Taxonomic profiling can then be performed either by region or via SIDLE reconstruction.
4. **IL paired-end known primers** This pipeline covers IL paired-end data, typically generated using primers targeting the V3–V4 region. Primer removal is carried out with cutadapt. Although most IL experiments target a single amplicon, the pipeline also supports cases with multiple amplicons, applying DADA2 per region when required. Taxonomic profiling is therefore usually region-based, but SIDLE reconstruction remains available when mixed amplicons are present.

After taxonomic assignment, an abundance table is generated containing the taxonomic classifications of features and their corresponding read counts per sample. Additionally, functional predictions using q2-metnet are supported.

#### Results Visualization and Output Generation

The final stage of the workflow involves the analysis of taxonomic and functional abundance tables and the automatic generation of summary figures, fully integrated into *Mbiome*. Abundance analysis is performed via complementary Python scripts and functional analysis figures are created by q2-metnet itself. In this way, *Mbiome* provides a complete workflow, from raw fastq input files to publication-ready figures and tables. Examples of these outputs are shown in Figures 3–7.

**Figure 3.**
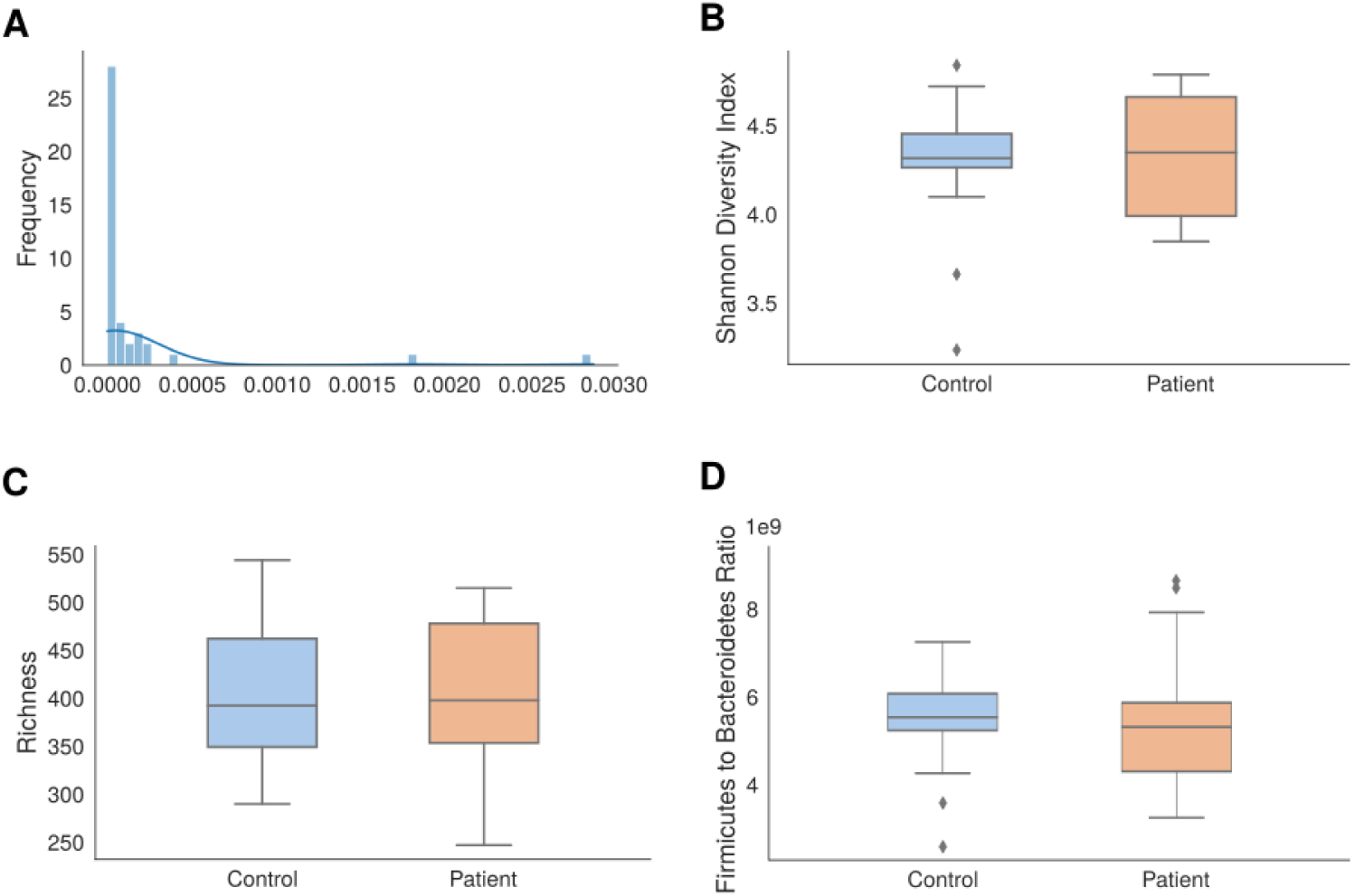
Overview of microbiome analysis outputs generated by the pipeline: (**A**) proportion of unclassified reads, (**B**) Shannon diversity index (SDI), (**C**) taxa richness, and (**D**) Firmicutes-to-Bacteroidetes (F/B) ratio, all stratified by sample type. These plots are automatically produced for any input dataset.

**Figure 4.**
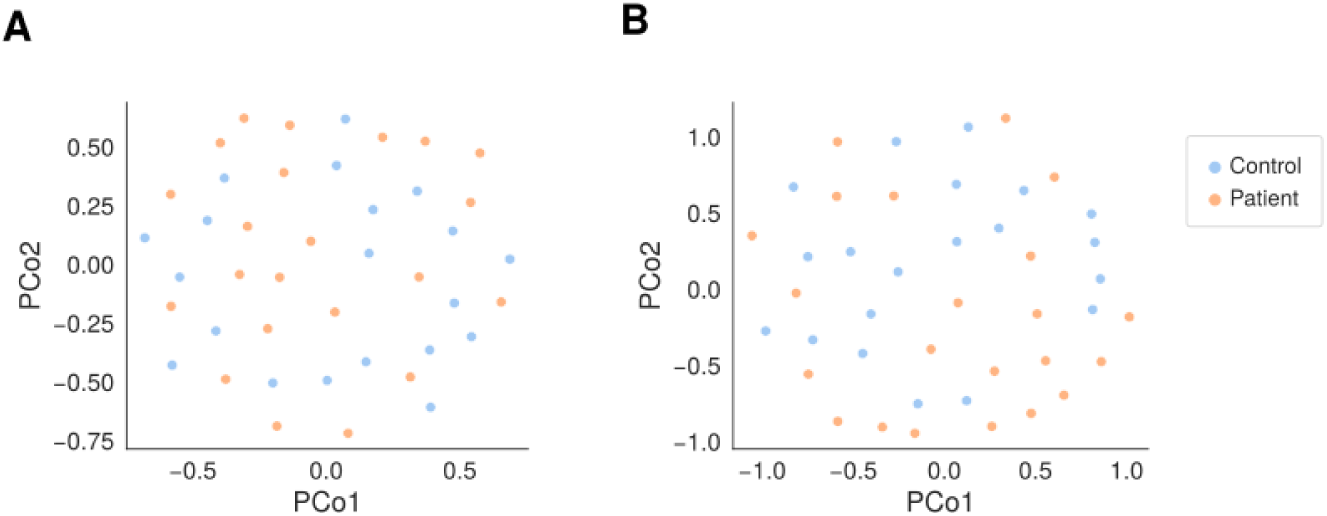
Overview of microbiome analysis outputs generated by the pipeline: PCoA plots of *β*-diversity (Bray-Curtis) using (**A**) precomputed distances and (**B**) euclidean.

**Figure 5.**
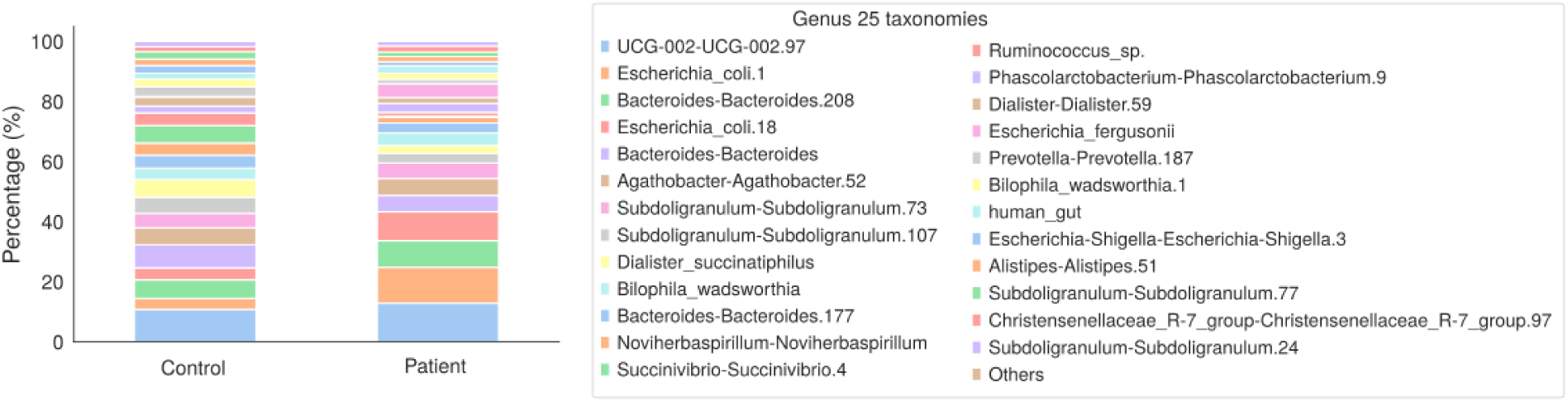
Examples of the Abundance analysis results: Relative abundance of the top 25 genera across groups.

**Figure 6.**
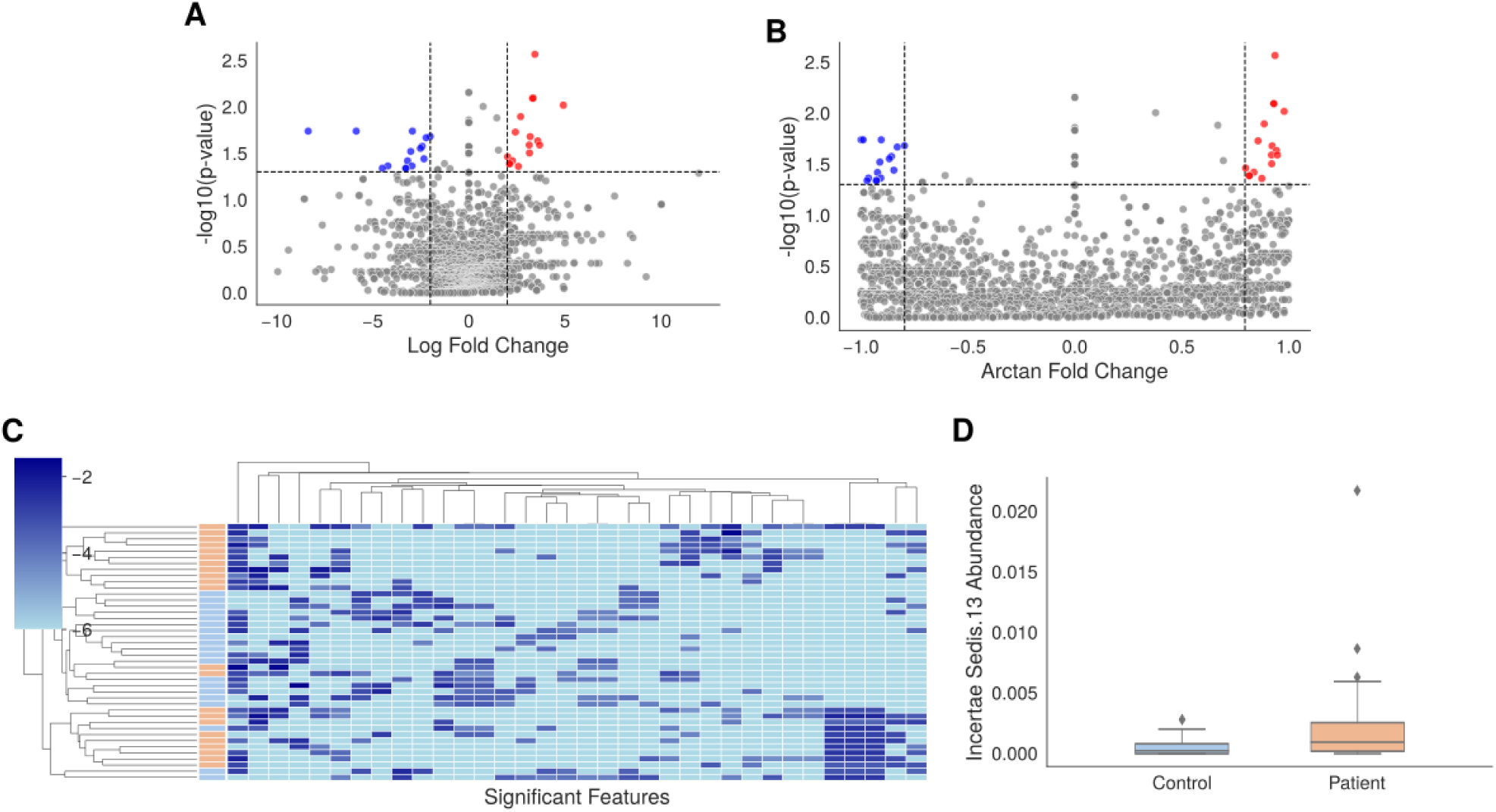
Overview of differential abundance analysis outputs generated by *Mbiome*: (**A**) volcano plot using Log Fold Change, (**B**) volcano plot using Arctan Fold Change — a bounded transformation that reduces the influence of extreme values —, (**C**) hierarchically clustered heatmap of statistically significant features across samples, and (**D**) example boxplot of one of the most significant taxon’s abundance across comparison groups. Red and blue dots indicate significantly up- and down-regulated taxa, respectively.

**Figure 7.**
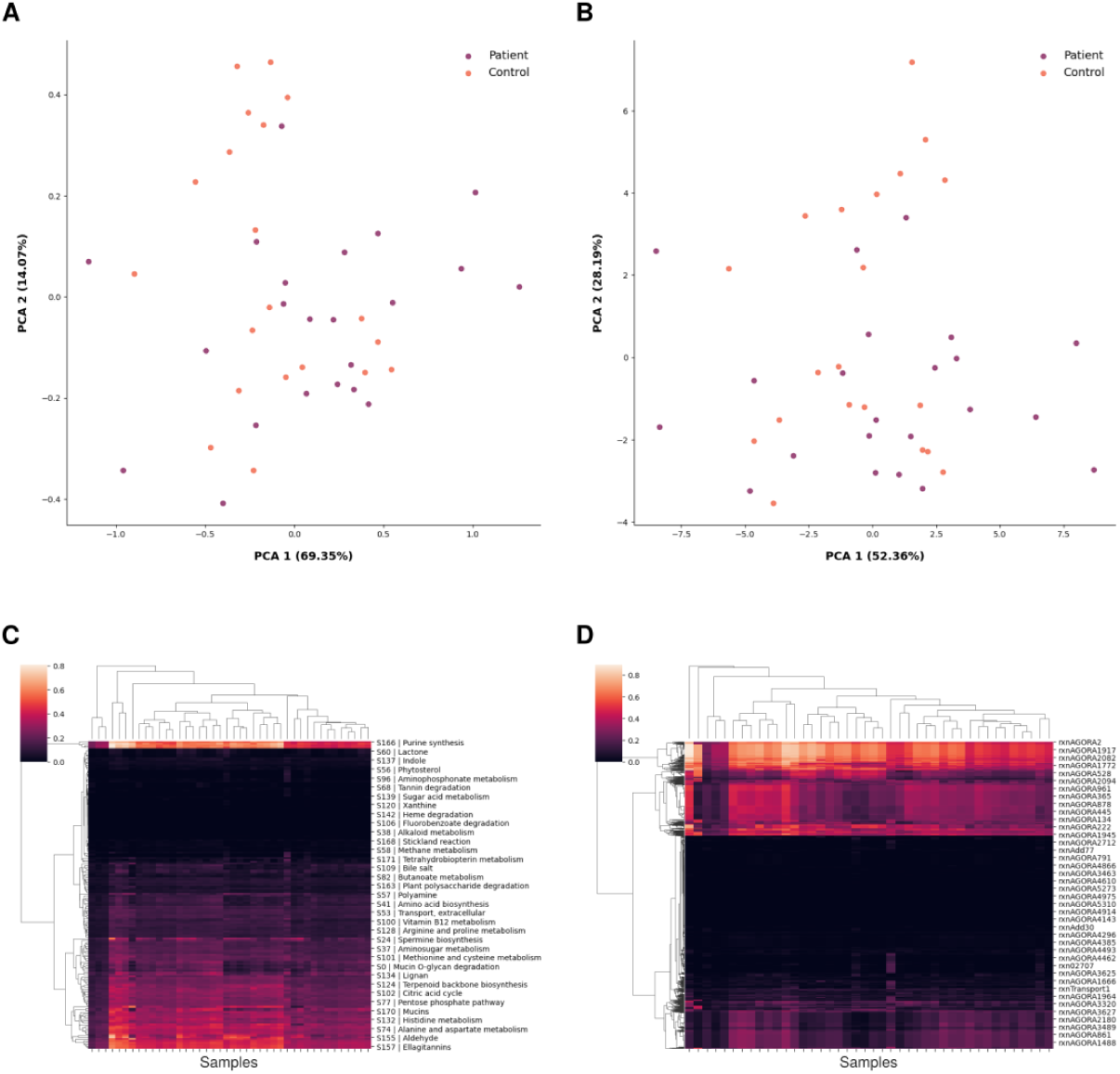
Overview of the q2-metnet functional analysis outputs: (**A**) PCA of metabolic subsystem activity scores, (**B**) PCA of individual reaction activity scores, both coloured by group membership (Patient/Control); (**C**) hierarchically clustered heatmap of metabolic subsystems, and (**D**) hierarchically clustered heatmap of individual reactions (AGORA identifiers). All outputs are generated automatically for any input dataset.

Figure 3A shows the distribution of unclassified reads across samples. These reads correspond to sequences that could not be assigned to any taxonomic group. Figures 3B and 3**C** present the Shannon diversity index (SDI) and richness, respectively. The SDI is an *α*-diversity metric that captures both richness and evenness within sample types, whereas richness refers only to the number of taxa detected. Together, these indices illustrate intra-group diversity and facilitate comparisons of microbial variation across metadata-defined categories. Another indicator of an overview of microbiota current state is the Firmicutes to Bacteroidetes (F/B) ratio, shown in Figure 3D, which reflects the balance between the two most dominant bacterial phyla in the gut microbiome. This ratio has been proposed as a proxy for gut dysbiosis, with elevated values associated with metabolic alterations such as obesity and insulin resistance, while lower values are generally linked to a healthier microbial composition (Ley et al., 2006; Turnbaugh et al., 2006; Magne et al., 2020). The boxplot illustrates the distribution of F/B ratios across the different sample groups, revealing both central tendency and variability within each category. Differences in the median and interquartile range across groups may suggest shifts in the overall community structure that are not captured by *α*-diversity metrics alone.

Regarding *β*-diversity, Figures 4A and 4**B** show Principal Coordinates Analysis (PCoA) plots based on Bray–Curtis dissimilarities, calculated using either precomputed distance matrices or Euclidean-transformed distances, respectively. Both visualizations illustrate the overall similarity structure of microbial communities across samples, highlighting clustering patterns according to their taxonomic composition.

Figure 5 displays the relative abundance of the 25 most prevalent genera, although the visualization can be adapted to any taxonomic level, providing insights into the dominant taxa in the dataset. *Mbiome* allows users to generate all these plots for any chosen metadata variable or groups.

A statistical analysis of the taxonomic information can also be performed through *Mbiome*. Figure 6 illustrates the differential abundance analysis outputs automatically generated by the pipeline. Figures 6A and 6**B** display volcano plots that combine effect size and statistical significance, using Log Fold Change and Arctan Fold Change, respectively. The latter provides a bounded transformation of the fold change that mitigates the visual distortion caused by extreme values, offering a complementary perspective on differential abundance. In both plots, taxa are coloured according to their statistical significance and direction of change: red dots indicate up-regulated taxa and blue dots indicate down-regulated taxa, while grey dots represent non-significant features. Taxon labels are omitted for visual clarity. Figure 6C presents a hierarchically clustered heatmap of all statistically significant features, enabling the identification of co-abundance patterns across samples; feature labels are omitted for clarity. Finally, Figure 6D shows a boxplot of one of the top differentially abundant taxon, automatically generated by *Mbiome* for each statistically significant feature — in this example, *Incertae Sedis* 13 — stratified by group, providing a direct visualisation of the abundance distribution underlying each individual statistical result.

Finally, the workflow also provides results about the functional analysis of the microbiome through the incorporated plugin (q2-metnet), which generates normalized metabolic activity scores for each reaction and subsystem, as well as differential statistical analyses between groups. Sample-level variation in metabolic potential is visualised through PCA plots at two levels of resolution: subsystem (Figure 7A) and individual reaction (Figure 7B), coloured by group membership. Complementarily, hierarchically clustered heatmaps display the metabolic activity profiles across samples for both subsystems (Figure 7C) and reactions (Figure 7D), enabling the identification of co-active metabolic modules and potential functional biomarkers differentiating the comparison groups.

### Evaluation of Sequencing Platforms and Analytical Pipelines

To assess the accuracy, robustness, and generalization of *Mbiome*, a comparative analysis involving different sequencing platforms and computational approaches was performed. This included *Mbiome* pipeline variants, as well as the external commercial solution, IR software. The goal was to understand how these factors affect taxonomic classification accuracy and microbiome diversity metrics.

In order to facilitate the understanding of these results, the nomenclature used for naming each pipeline and a description of each of them is described in Table 4 (in Methods section).

#### Total reads obtained per sample differ by method

Figure 8 shows the total reads per samples obtained by each method. It is noteworthy that when using IL sequencing platform, the total count of reads per sample shows much higher variation, followed by ITkitNoSplit. The rest of the methods, however, show a remarkably similar total amount of reads per sample, coincidentally, the three methods that use the splitting by region approach for the analysis. This might be due to a normalization when performing the reconstruction of regions.

**Figure 8.**
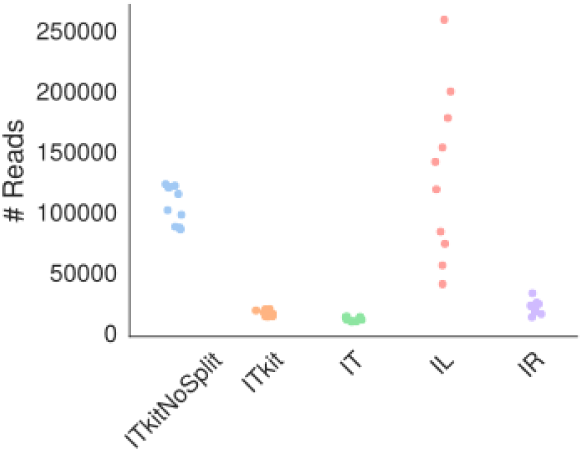
Total read count at genus taxonomic level per sample by method.

#### Mbiome achieves superior genus-level classification compared to commercial software

The taxonomic classification efficiency at the genus level was evaluated. In microbial community profiling, some reads cannot be confidently assigned down to the genus rank due to limitations in sequence quality, reference databases, or inherent ambiguity. These are referred to as unclassified reads at the genus level. Figure 9A shows the proportion of unclassified reads for each method. Although the commercial IR pipeline does not display unclassified reads in its main output—since they are stored in a separate file—it ultimately classified the smallest fraction of reads at the genus level, meaning a larger share of reads remained unresolved at this taxonomic rank.

**Figure 9.**
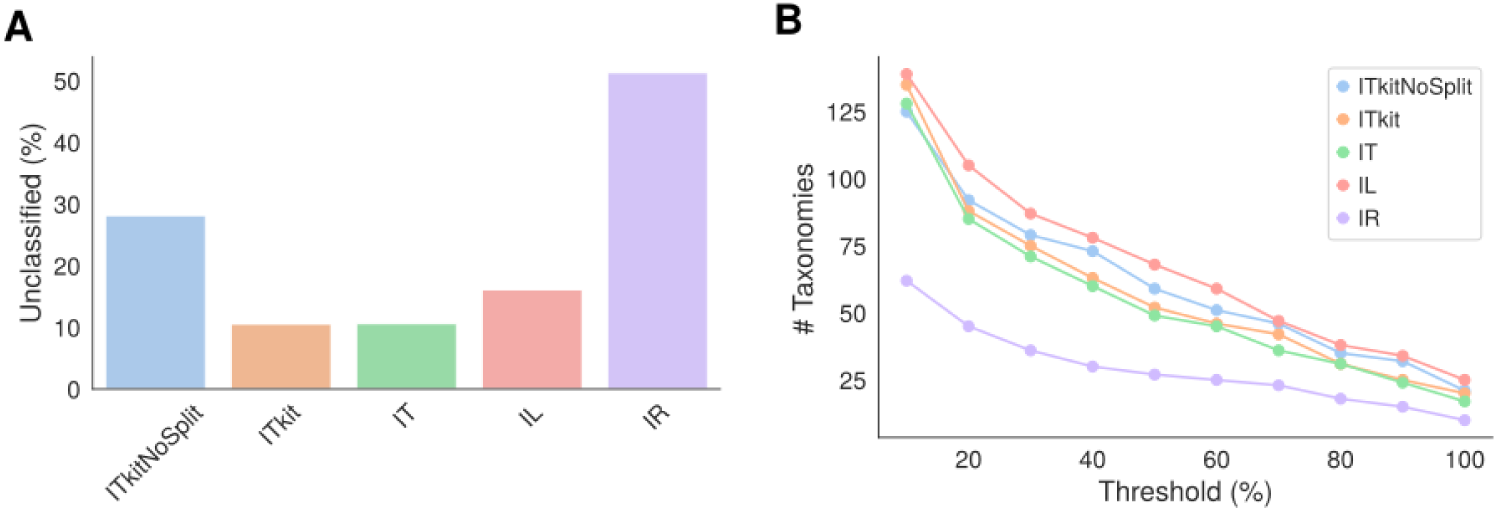
Comparison of pipeline performance across methods based on percentage of unclassified reads and unique taxonomies at the genus level: **(A)** Percentage of unclassified reads at genus taxonomic level by method. **(B)** Number of unique taxonomies for each method by different filtering thresholds.

Figure 9B compares the total number of unique genus-level taxa detected by each method across varying abundance thresholds. Methods without region splitting (ITkitNoSplit and IL) identified the greatest number of unique taxa, followed by the IT-based *Mbiome* pipelines. The IR method reported the fewest unique taxa, likely reflecting pre-processing steps that eliminate unclassified taxa and assign reads to the deepest taxonomic rank.

#### Mbiome pipelines show higher consistency in genus-level classification than commercial software

To evaluate the consistency of taxon detection across pipelines, the overlap of genus-level classifications among methods was analyzed. Different prevalence thresholds were applied, requiring taxa to appear in at least 10%, 20%, or 30% of samples. As shown in Figure 10, even under lenient filtering, approximately 30–35% of genera were shared by four out of the five methods.

**Figure 10.**
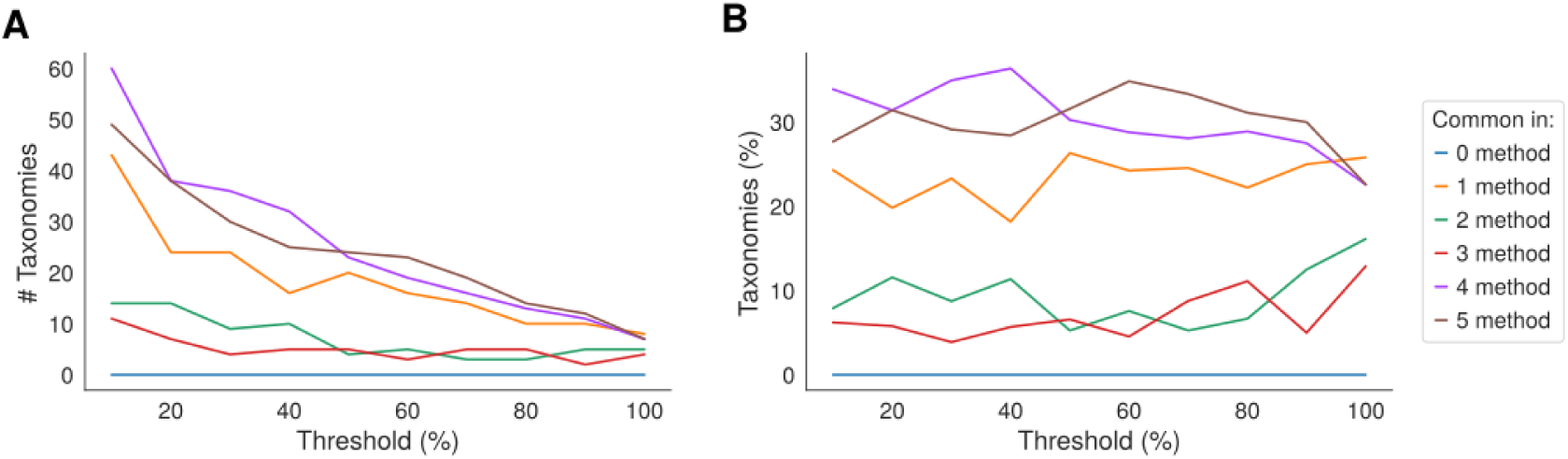
Number of genera shared across a given number of methods asa function of the prevalence filtering threshold, expressed in absolute counts (**A**) and as a percentage of the total detected genera (**B**). Each line represents the subset of taxa detected in exactly 0 to 5 methods simultaneously.

To further explore how these taxa are distributed across methods, Figure 11 illustrates the intersections of shared genera under the three prevalence thresholds (10%, 20%, and 30%). Across all thresholds, the largest intersection consistently corresponds to taxa shared by all five pipelines simultaneously, indicating a high degree of consensus among the methods. The IR workflow, which differs the most in its processing and analytical approach, is the one most frequently absent from the core consensus group. In contrast, the IL method shows the highest number of unique taxa.

**Figure 11.**
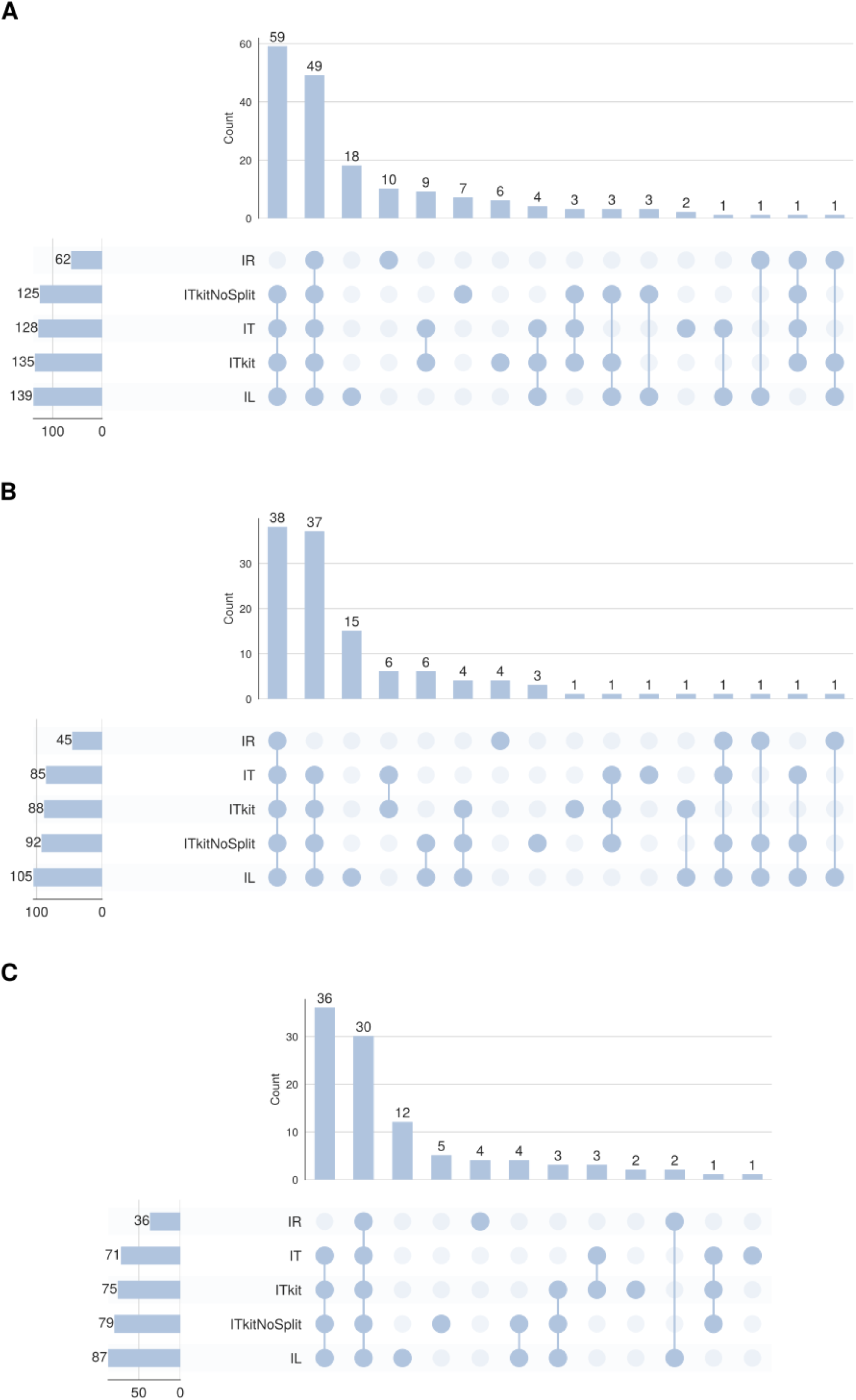
UpSet plots showing the intersections of genus-level taxa detected by each pipeline (IR, IT, ITkit, ITkitNoSplit, IL) at prevalence thresholds of (**A**) 10%, (**B**) 20%, and (**C**) 30%. Bar height indicates the number of genera in each intersection; horizontal bars on the left show the total number of genera detected per method.

The taxonomic similarity was also quantified, using the Jaccard index (Figure 12). The three IT *Mbiome* pipelines—ITkit, IT, and ITkitNoSplit—exhibited the highest similarity scores across all prevalence thresholds, reflecting consistency within the software framework despite differences in splitting strategy. In contrast, the commercial IR pipeline consistently showed the lowest similarity to all other methods, reinforcing its distinct classification profile. Jaccard scores between non-IR pipelines generally ranged from 0.70 to 0.90, suggesting broadly comparable genus-level profiles. Comparisons involving IR rarely exceeded 0.50, further highlighting its divergence from the remaining pipelines.

**Figure 12.**
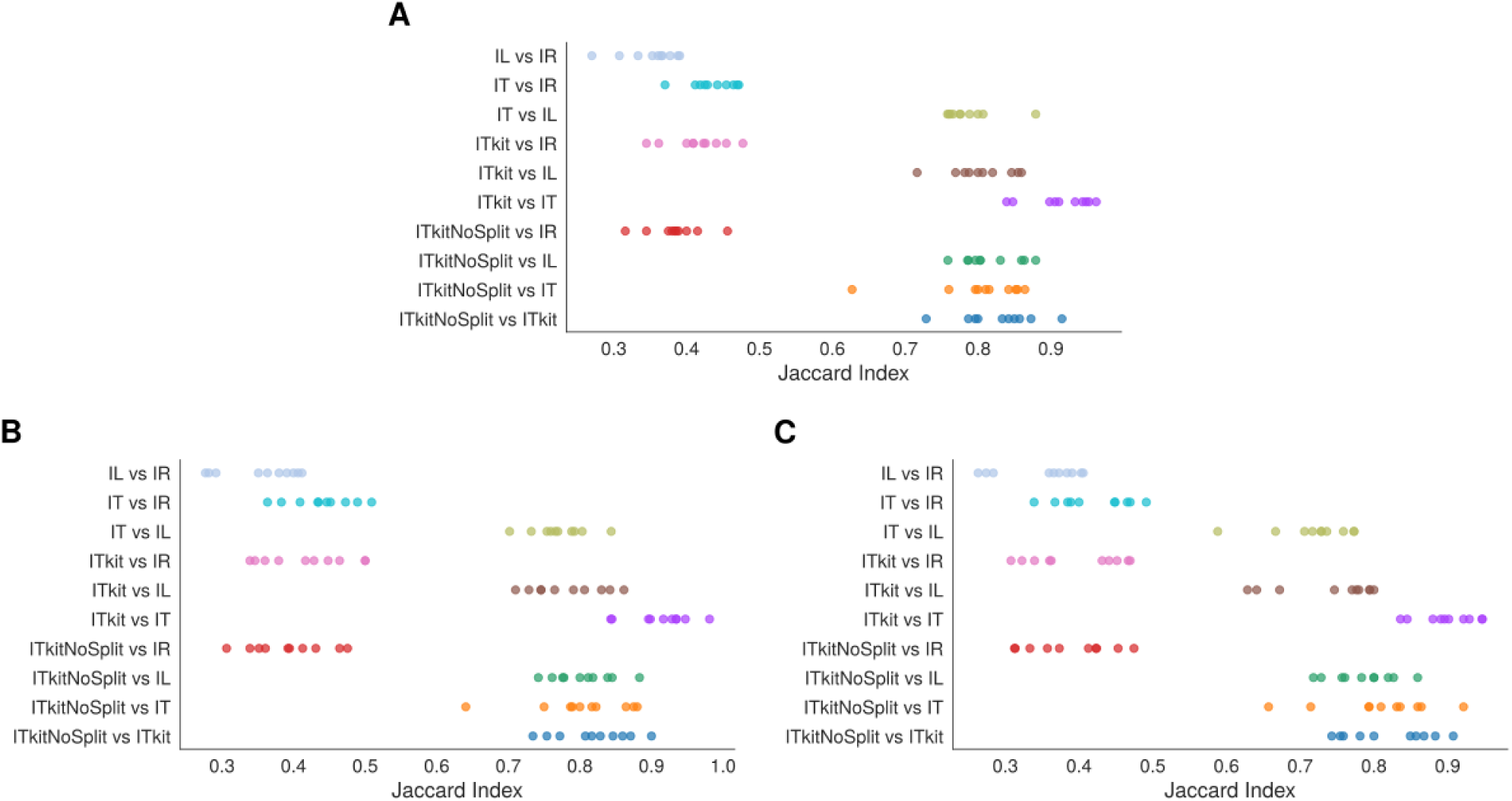
Pairwise Jaccard similarity indices between all method pairs at prevalence thresholds of 10% (**A**), 20% (**B**), and 30% (**C**). Each dot represents a sample-level comparison; the x-axis indicates the proportion of shared genera between each pair of methods relative to their union.

#### Genus-level diversity and richness differ across classification methods

To complement taxonomic presence analyses, ecological diversity metrics at the genus level were compared using a 30% prevalence threshold. This threshold was selected to balance the inclusion of common taxa while maintaining robustness given the modest sample size (10 samples). Filtered abundance tables were used to focus comparisons on taxa consistently detected across methods.

The SDI and richness were calculated for each method; statistical results are provided in Tables 2 and 3. As shown in Figure 13**A**, SDI values were broadly comparable across the IT-based pipelines (ITkitNoSplit, ITkit, IT) and IL, with medians ranging approximately from 2.6 to 3.1. IR, however, showed markedly lower SDI values (median *≈* 2.0), further corroborating its distinct classification profile. ITkitNoSplit also displayed slightly lower SDI compared to the remaining IT-based methods.

**Figure 13.**
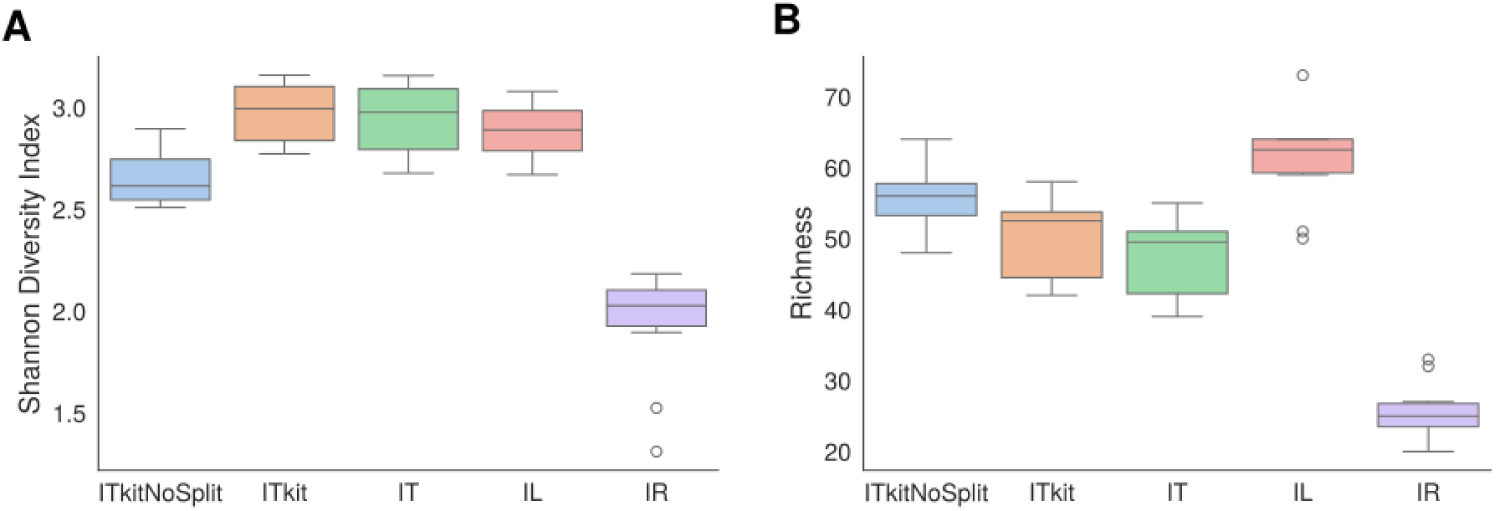
Comparison of genus-level ecological diversity metrics across classification methods at a 30% prevalence threshold: (**A**) Shannon Diversity Index and (**B**) taxon richness. Boxes represent the interquartile range, horizontal lines indicate the median, and open circles denote outliers.

**Figure 14.**
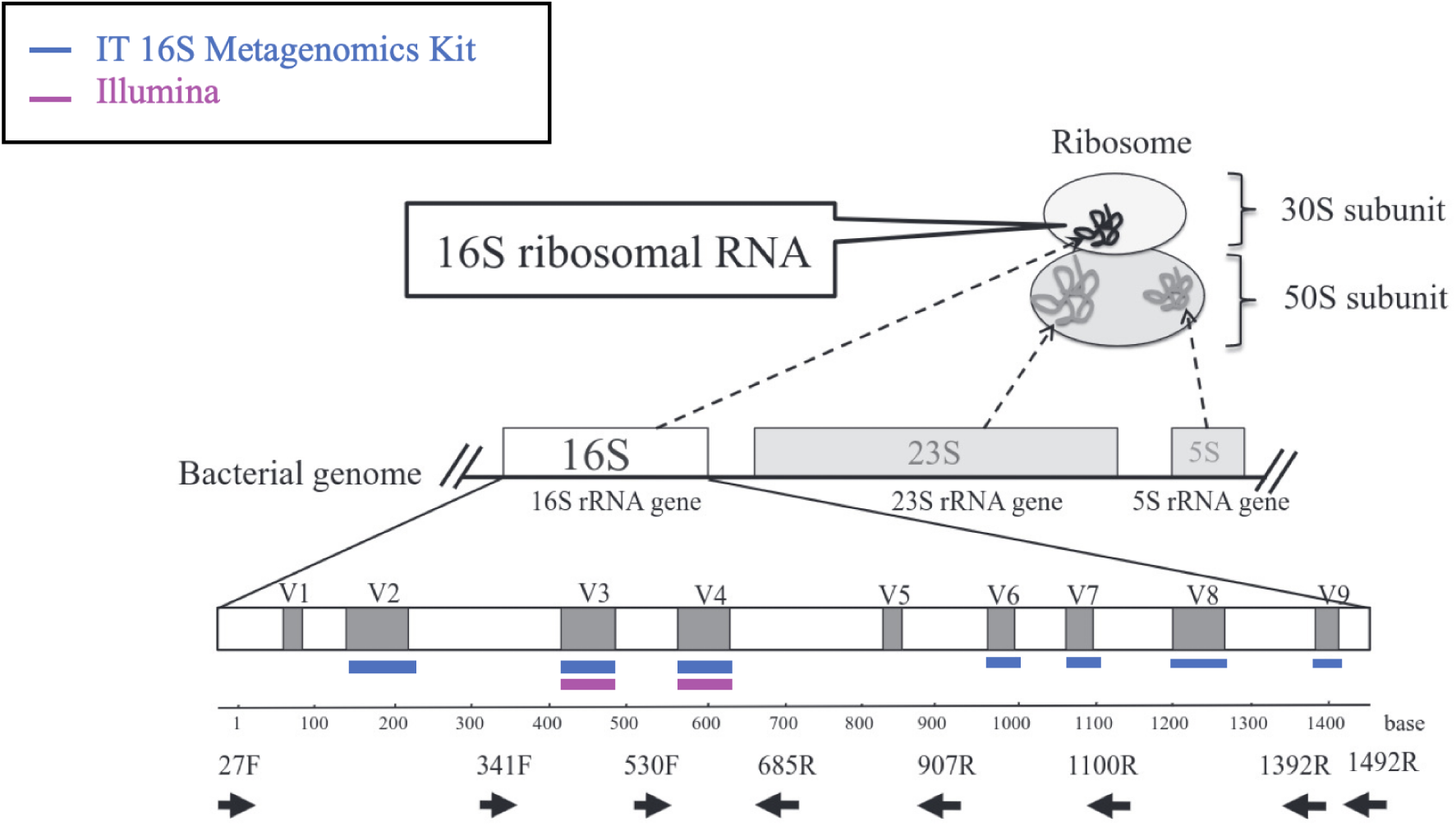
Differences between IT and IL in the amplification approach of different hypervariable regions.

**Table 2.** P-values for pairwise comparisons of SDI at genus level and threshold 30.

| Method 1 | Method 2 | P-value |
| --- | --- | --- |
| ITkitNoSplit | ITkit | **0.000769** |
| ITkitNoSplit | IT | **0.002202** |
| ITkitNoSplit | IL | **0.003611** |
| ITkitNoSplit | IR | **0.000183** |
| ITkit | IT | 0.520523 |
| ITkit | IL | 0.161972 |
| ITkit | IR | **0.000183** |
| IT | IL | 0.273036 |
| IT | IR | **0.000183** |
| IL | IR | **0.000183** |

**Table 3.** P-values for pairwise comparisons of Richness at genus level and threshold 30.

| Method 1 | Method 2 | P-value |
| --- | --- | --- |
| ITkitNoSplit | ITkit | 0.095177 |
| ITkitNoSplit | IT | **0.007984** |
| ITkitNoSplit | IL | **0.044751** |
| ITkitNoSplit | IR | **0.000179** |
| ITkit | IT | 0.225076 |
| ITkit | IL | **0.005026** |
| ITkit | IR | **0.000178** |
| IT | IL | **0.001293** |
| IT | IR | **0.000180** |
| IL | IR | **0.000180** |

Richness differences were more pronounced (Figure 13**B**) consistent with the significant p-values reported in Table 3. IR exhibited the lowest richness (median *≈* 25 genera), whereas IL showed the highest values (median *≈* 60 genera), consistent with the broader detection profile observed in the UpSet analysis (Figure 11).

## Discussion

The development of an automated workflow for microbiome analysis represents an important step toward the democratization of bioinformatics, enabling a broader community of researchers and clinicians to perform reproducible and transparent analyses. By integrating QIIME2 as the core framework, *Mbiome* combines robustness and accessibility, eliminating the need for programming or command-line expertise. This effort aligns with the increasing demand for open, standardized, and reproducible computational tools, particularly as microbiome research expands into clinical and translational applications (Xia and Sun, 2023; Rojas-Velazquez et al., 2024).

### Strengths and contributions of Mbiome

The comparative evaluation demonstrated that analyses conducted with *Mbiome* achieved more consistent and robust taxonomic profiles than those obtained using the IR commercial software, which is commonly employed for Ion Torrent data. Although *Mbiome* is built upon the QIIME2 framework, its automation of all necessary steps and parameter optimization enhances usability and reduces human error, while maintaining transparency and flexibility.

The full automation of microbiome analysis—from raw fastq files to final abundance tables, functional predictions, and visual outputs—greatly facilitates accessibility and interpretability. Moreover, the inclusion of metabolic pathway inference provides a more comprehensive understanding of microbial communities, highlighting the microbiome’s functional role beyond taxonomic composition. This systems-level perspective is essential for elucidating microbiota-driven processes in health and disease (Dakal et al., 2025).

Another key advantage of *Mbiome* is its flexibility: it supports data from multiple sequencing platforms (Illumina and Ion Torrent), different molecular targets (16S and ITS), and various primer designs, including kit-specific approaches. Such versatility makes it suitable for a wide range of research contexts, from clinical studies to environmental microbiology, while ensuring methodological rigor through optimized parameters for each step (Maki et al., 2023).

### Impact of methodological choices on results

Application of *Mbiome* to real datasets revealed that methodological variations in sample processing and bioinformatic pipelines can markedly affect the outcomes of microbiome analyses. Even in the absence of direct biological comparisons, the observed discrepancies in the number of unique taxa or in the percentage of unclassified reads highlight how methodological inconsistency can lead to divergent interpretations. These findings reinforce the importance of standardizing workflows, documenting analytical decisions, and ensuring reproducibility—principles central to the FAIR data movement. *Mbiome* directly addresses these challenges by providing a transparent, traceable, and user-guided framework. This aligns with recent efforts to improve reproducibility in microbiome research through transparent and well-documented pipelines (Rojas-Velazquez et al., 2024).

### Transparency and user empowerment

Microbiome studies frequently rely on highly automated “black-box” software, which obscures analytical decisions and hinders reproducibility. *Mbiome* overcomes this limitation by combining automation with user control. Users can visualize quality plots, adjust parameters interactively, and select among integrated methodological options, fostering a deeper understanding of how analytical choices influence results. This user-centered approach encourages more critical and informed data interpretation.

### Limitations and future perspectives

While *Mbiome* provides a comprehensive and reproducible solution for amplicon-based microbiome analysis, the field itself is rapidly evolving. Emerging standards such as the nf-core pipelines—particularly nf-core/ampliseq—offer modular, scalable, and community-curated workflows built on Nextflow DSL2. These pipelines share many design principles with *Mbiome*, including support for multiple sequencing platforms (Illumina, Ion Torrent, PacBio), diverse molecular targets (16S, ITS, CO1, 18S), and integration with QIIME2 for taxonomic profiling and diversity analysis (Straub et al., 2020). Future developments of *Mbiome* will focus on interoperability with these frameworks, expansion to shotgun metagenomics and metabolomics, and incorporation of machine learning modules for predictive and integrative analysis (Langer et al., 2025)

In summary, *Mbiome* contributes to the ongoing effort to make microbiome research more accessible, standardized, and transparent. Its open-source release on GitHub (https://github.com/MGorostidi/mbiome) provides a platform for continuous improvement through community feedback and collaboration, ensuring that it remains aligned with the evolving standards of reproducible bioinformatics.

## Methods

### MBiome

The workflow encompasses the subsequent steps:

#### Requirements

In order to run a experiment, it is necessary to prepare three main files (*Mbiome* includes a EXAMPLE FILES folder with all the necessary information):

- samples-manifest.tsv file: as specified by QIIME2, this file is necessary in order to import fastq files. Depending on the characteristics of the experiment and the previous sample processing, this file will include different columns. Since samples can be processed in different sequencing runs, a manifest file for each run will be necessary and these should be renamed as “samples-manifest-runXXX.tsv”, being XXX a number from 1 to X (example: if only 1 run was performed, samples-manifest-run1.tsv will be enough, but if 2 sequencing runs were necessary, samples-manifest-run1.tsv and samples-manifest-run2.tsv will be created).
- samples-metadata.tsv file: this file collects all the metadata information about the samples. Whatever information can be added, but the format is important (check EXAMPLE FILES folder). A unique file will contain all the information about all the samples, even if different sequencing runs are present.
- config.env file: the configuration file that contains all the information about the characteristics of the experiment (already described in Table 1).

#### Data input

*Mbiome* accepts demultiplexed fastq files from IT or IL sequencing platforms, either from 16S Metagenomic Kit or specific primer sequences DNA amplification. Data is imported using samples-manifest.tsv file, as specified by QIIME2.

#### Quality control

If primer sequences are known, primer removal is performed using cutadapt (Martin, 2011). For the “IT known primers” pipeline, the cutadapt trim-single command is applied with parameters optimized for 5’ primer detection (e.g., –p-front with the “^” anchor, as recommended in (Maki et al., 2023)), and reverse primer trimming is enabled through –p-adapter. Additional options such as –p-indels and –p-error-rate 0.1 are included to improve mismatch tolerance during adapter detection, together with wildcard handling and untrimmed-read discarding (Qiime2-developers and Gorostidi-Aicua, 2022b). For the “IL paired-end known primers” pipeline, the same parameters are applied using the cutadapt trim-paired command, which requires explicit specification of forward and reverse primers (–p-front-f, –p-front-r). In the “IT Metagenomics Kit (byVregion)” pipeline, primer removal and region splitting are conducted using the proprietary MetagenomicsPP plugin (ThermoFisher), as described in Maki et al. (2023). Finally, in the “IT Metagenomics Kit” pipeline without primer information, primer sequences are implicitly handled during denoising with DADA2 (explained in Denoising sequences section).

#### Denoising sequences

Sequence quality is first assessed using the demux plugin in QIIME2, and reads are subsequently processed with DADA2 (Callahan et al., 2016). This step includes read trimming (when required), error correction, quality-based truncation, merging of paired-end reads, and chimera removal, resulting in the generation of amplicon sequence variants (ASVs). The quality-based truncation step is interactively determined by the user from the demultiplexed quality plots generated during pipeline execution, following the criteria described in the workflow.

For pipelines with known primers, as well as for the “IT Metagenomics Kit (byVregion)” pipeline, only the –p-trunc-len parameter is applied, since primer sequences has already been removed. In IL paired-end datasets, forward and reverse reads are truncated to the same length to ensure compatibility. This truncation step removes low-quality bases from read tails, ensuring uniform sequence length.

In the “IT Metagenomics Kit” pipeline, an additional trimming of 15 bp is performed with –p-trim-left, following IT specific recommendations (Qiime2-developers and Jones, 2020; Qiime2-developers and Gorostidi-Aicua, 2020).

When experiments include multiple sequencing runs, DADA2 is executed independently per run, and feature tables and representative sequences are merged afterwards (Qiime2-developers and Gorostidi-Aicua, 2022a). Prior to merging, denoising statistics are evaluated by the user to confirm adequate quality; if necessary, truncation parameters are adjusted and the denoising step is repeated.

#### Taxonomic assignment

Representative sequences were taxonomically classified using the qiime feature-classifier classify-consensus-vsearch plugin implemented in QIIME2 (Rognes et al., 2016). For 16S rRNA gene data, either the SILVA (Quast et al., 2012) or Greengenes (DeSantis et al., 2006) reference databases were employed, while the UNITE database (Nilsson et al., 2018) was used for ITS sequences.

Sequences were aligned against the selected reference database using local similarity searches, and taxonomy was assigned according to the consensus of the best hits. Unless otherwise specified, default QIIME2 parameters were applied, including a minimum sequence identity of 80% and a minimum query coverage of 80%, following recommended guidelines. In cases where multiple hypervariable regions of the 16S rRNA gene were sequenced separately, taxonomic profiles were reconstructed using the SIDLE (SMURF Implementation Done to acceLerate Efficiency) reconstruction, through q2-sidle plugin (Debelius, 2020). SIDLE is a python version of the Short MUltiple Reads Framework (SMURF) algorithm (Fuks et al., 2018) with a novel tree-building approach, that allows the reconstruction of multiple short, fragmented amplicons against a known database to improve the resolution of the reconstructed community over single amplicons (Debelius, 2020).

#### Metabolic pathway inference

To infer the metabolic potential of microbial communities from 16S rRNA gene sequencing data, the q2-metnet plugin was used within the QIIME2 framework (Balzerani et al., 2024). This tool contextualizes taxonomic profiles into curated metabolic reconstructions, specifically AGORA and AGREDA, enabling functional interpretation of microbial composition. Based on the relative abundances of taxa, q2-metnet computes normalized activity scores for metabolic reactions and subsystems. These scores were used to perform differential activity analyses and exploratory visualizations, including PCA and hierarchical clustering, to identify functional differences between sample groups.

#### Abundance table analysis

Microbial diversity was evaluated through both *α*- and *β*-diversity metrics. *α*-diversity was quantified using the SDI, which incorporates richness and evenness, and by genus richness, defined as the number of taxa with non-zero abundance per sample. The SDI was computed from relative abundances obtained by normalizing taxon counts and applying the Shannon entropy formula (SDI and richness are explained in Evaluation metrics section). *β*-diversity was assessed using pairwise distance matrices, primarily the Bray–Curtis dissimilarity metric, which captures compositional differences in community structure between samples.

#### Statistical analysis

Group comparisons were performed based on data distribution and the number of groups defined by the dependent variable. Normality within groups was assessed using the Shapiro–Wilk test. For normally distributed data, comparisons between two groups were conducted using a two-sample t-test, while comparisons involving more than two groups were performed using one-way ANOVA. For non-normally distributed data, the Mann–Whitney U test (two groups) and the Kruskal–Wallis H test (more than two groups) were applied. Group sizes and total sample counts were documented for each analysis. To correct for multiple testing, p-values were adjusted using the Benjamini–Hochberg false discovery rate (FDR) procedure.

#### Visualization and reporting

Summary figures and tables were generated in Python using matplotlib and seaborn for taxonomic barplots, boxplots, principal coordinates analysis (PCoA) plots, and comparative analyses. UpSet diagrams were produced with the upsetplot package.

#### Reproducibility

The complete workflow is implemented as a Bash and Python pipeline under version control. All scripts and configuration files are publicly available at https://github.com/MGorostidi/mbiome.

### Evaluation of Sequencing Platforms and Analysis Approaches

To evaluate potential biases introduced by sequencing technologies and analysis pipelines, a comparative analysis was performed using two widely adopted platforms: Illumina (IL) and Ion Torrent (IT). Given the predominant use of IL in microbiome research, this comparison aimed to determine the extent to which platform choice influences downstream microbial community profiling.

Ten stool samples were selected for this study. Each sample underwent DNA extraction followed by sequencing on both IT and IL platforms, thereby minimizing biological variability and allowing differences to be primarily attributed to sequencing methodology.

Five distinct analytical pipelines were included in the comparison to evaluate how different processing strategies influence microbiome profiling outcomes. These comprised four approaches provided within the *Mbiome* framework and one commercial software, Ion Reporter (IR), a commercial software provided by Thermo Fisher Scientific. The nomenclature used for each pipeline and a description of each of them is included in Table 4.

**Table 4.** Summary of pipelines for 16S metagenomics analysis.

| Pipeline | Abbreviation | DNA amplification | Platform | Computational approach |
| --- | --- | --- | --- | --- |
| IT Metagenomics Kit | ITkitNoSplit | 16S Metagenomics Kit | Ion Torrent | No splitting by V region. Trimming of 15 bp during DADA2 for primer removal. |
| IT Metagenomics Kit by V region | ITkit | 16S Metagenomics Kit | Ion Torrent | Splitting by V region using MetagenomicsPP plugin. Analysis by region and taxonomic reconstruction by SIDLE. |
| IT known primers | IT | Commercial primers | Ion Torrent | Splitting by V region and primer removal by cutadapt. Analysis by region and taxonomic reconstruction by SIDLE. |
| IL paired-end known primers | IL | Commercial primers | Illumina | Standard Illumina paired-end analysis pipeline. *The rest of methods include different V regions (V2, V3, V4, V6-7, V8 and V9), while this pipeline only a unique amplicon of V3V4. |
| Ion Reporter | IR | 16S Metagenomics Kit | Ion Torrent | Analysis using IR software suite. The software splits by region and then a consensus taxonomy abundance table of all regions is calculated. |

The commercial IR pipeline was included as an external benchmark, representing a widely used proprietary solution. To ensure a fair and consistent comparison, the IT-based pipelines (*IR*, *ITkit*, *ITkitNoSplit*, and *IT*) were all applied to the IT sequencing data, whereas the *IL* pipeline was applied exclusively to IL data. The same ten stool samples were used throughout, with identical DNA extracts sequenced and analyzed independently by each method. Even *Mbiome* includes both SILVA and GreenGenes as optional reference databases for the taxonomic classification of the bacterial community, for this comparative analysis SILVA was used.

This experimental design enabled a systematic assessment of how sequencing platform and computational analysis interact to affect taxonomic classification and microbial diversity metrics.

#### Data Processing

Raw sequencing reads from both IT and IL platforms were processed through their respective analysis pipelines as described. IT platform data were analyzed using the IR pipeline and three *Mbiome* IT pipelines, while IL data were processed using the *Mbiome* IL pipeline.

Taxonomic feature names were systematically standardized by extracting primary taxonomic terms across ranks (domain to genus) and removing extraneous characters such as underscores, hyphens, and bracketed annotations. This step ensured uniformity across datasets and facilitated downstream comparisons.

Duplicate taxonomic features identified by identical names were merged by summing their abundance values to eliminate redundancy. Features lacking assignment at the genus level (comparative analysis was performed at this taxonomic level) were labeled as “Unclassified”.

To harmonize taxonomic identifiers, cleaned taxonomic names were queried against the NCBI Taxonomy database (Federhen, 2011) using the Entrez API to retrieve corresponding Taxonomy IDs (TaxIDs). Features without unique TaxID matches were excluded. Finally, columns sharing the same TaxID were consolidated by summing their abundances, resulting in a harmonized abundance matrix suitable for comparative analysis.

#### Evaluation Metrics

The following metrics were employed to compare sequencing platforms and analysis pipelines:

- **Total Reads per Sample:** Assessment of total sequencing reads obtained per sample across methods to evaluate consistency and variability in sequencing depth.
- **Unique Taxonomies:** Quantification of unique taxonomic units detected at the genus level by each pipeline to assess taxonomic resolution and sensitivity.
- **Taxonomic Overlap:** Analysis of shared and unique taxonomies among methods using upset plots and Venn diagrams. Taxa prevalence thresholds of 10%, 20%, and 30% of samples were applied to focus comparisons on commonly detected taxa.
- **Jaccard Similarity Index:** A statistical measure of similarity between taxonomic sets identified by different methods based on presence/absence, defined as the size of the intersection divided by the size of the union of the two sets. It ranges from 0 (no overlap) to 1 (identical sets), and is calculated as:

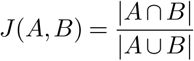

where *A* and *B* represent the sets of taxonomies identified by two different methods.
- **Ecological Diversity Metrics:** SDI and Richness were calculated at the genus level to assess microbial community diversity and complexity across methods.

**– Richness:** Defined as the total number of distinct taxa (e.g., genera) observed in a sample. It reflects the breadth of the community but does not account for relative abundances.
**– Shannon Diversity Index (SDI):** A measure that accounts for both richness and evenness of taxa in a community. It is calculated as:

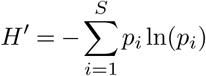

where *S* is the total number of taxa and *p_i_* is the relative abundance of taxon *i*. Higher values indicate more diverse and evenly distributed communities.

### Sample collection and processing

#### Sample collection

Stool samples of people with Multiple Sclerosis (pwMS) from the *Hospital Universitario Donostia* (Donostia-San Sebastián, Spain) were collected. Written informed consent was obtained for each participant before inclusion. All the samples had been recruited under projects accepted by the Ethical committee from our institution.

All participants were provided with a comprehensive kit containing all the necessary materials for hygienic fecal sampling in the comfort of their own home. The kit included a stool collector designed to adhere to the toilet, tubes to collect stool samples, a safety bag, and hydrated cold packs to maintain the required temperature. An isothermal bag was also provided for safe transport of the samples.

Upon collection, participants deposited their stool sample into the provided tube, which was immediately frozen at *−*20 *^◦^*C. The samples were then transported to the hospital, using the cold packs included in the kit. On arrival at our center, the samples were thoroughly checked for suitability before being stored at *−*80 *^◦^*C until further analysis.

#### DNA extraction and sample processing

Fecal samples were thawed on ice and diluted in 1X Dulbecco’s phosphate-buffered saline (DPBS) (Gibco BRL, Gaithersburg, MD, USA). DNA was then extracted through the mechanical and enzymatic lysis of the cells and the extraction kit QIAamp DNA Stool Mini Kit (Qiagen, Hilden, Germany). Mechanical disruption was performed by bead-beating with 0.1 mm diameter zirconia/silica beads three times (Sigma, St. Louis, MO, USA) using a Bead Ruptor 12 (OMNI International, Georgia, USA). Enzymatic lysis of fungal cells was performed using the enzyme zymolyase (MP Biomedicals, LLC, Illkirch-Graffenstaden, France) at a concentration of 0.1 mg/mL. The enzyme lysozyme (Sigma, St. Louis, MO, USA) at a concentration of 10 mg/mL was used for the bacterial lysis. The obtained DNA was subjected to quality and quantity controls.

#### Sample sequencing

Both IT and IL sequencing equipment were used in order to test the performance of all the possible workflows included in *Mbiome*, comparing both platforms in order to evaluate the bias of the sequencing tool in the results.

Apart from the technical differences between sequencing methods used by IT and IL, such as, the sequencing method, time, read quality and length, or cost, other key dissimilarities exist, that should be taken into account for posterior analyses. For this project, the principal difference between the sequencing approaches is the number of hypervariable regions included in the amplification and the sequencing steps.

The number of hyervariable V regions amplified, thus, the amount of amplicons produced by each sequencing method, varies (Figure 14). When running the analysis, it is important to know if the samples contain reads proceeding from from different amplicons or not, in order to perform one approach or the other. For this project, IT data commonly contained a mix of different amplicons, due to the Ion 16S Metagenomics Kit (by Thermofisher) being used. The kit amplifies 7 out of the 9 hypervariable regions (specifically, V2, V3, V4, V6-7, V8 and V9). Since the amplification step is performed creating different amplicons, it is necessary to conduct the taxonomical analysis in a different way than it is done when a unique amplicon is created, as in the case of the IL part of this project, where a single amplification of V3 and V4 hypervariable regions is performed.

On the other hand, while IL sequencing can be single or paired-end, the second one producing two different fastq files containing forward or reverse sequences, IT does not contain that option. IT data contain mixed-orientation reads in the same file, what should be taken into consideration when choosing different parameters for running the algorithm.

The wet-lab protocols followed for each sequencing method are explained in the subsequent subsections:

##### Ion Torrent sequencing

For the IT equipment (Life Technologies, MA, USA) sequencing method between 300 and 400 ng of DNA was used to perform the amplification and sequencing.

The Ion 16S Metagenomics Kit (Thermo Fisher, Waltham, MA, USA) was used to amplify bacterial 16S rRNA V2, V3, V4, V67, V8, and V9 hypervariable regions. Barcode libraries were created using the Ion Plus Fragment Library Kit (Thermo Fisher, Waltham, MA, USA). The final library concentration was quantified (Kapa Library Quantification Kit (Roche, Basel, Switzerland)), and 26 pM of DNA was loaded onto IT 318 chips.

For fungal kingdom analysis, amplification of the fungal ITS1 and ITS2 intergenic regions was carried out using the following primers: ITS1 30F (5’-GTCCCTGCCCTTTGTACACA-’3) and ITS1 217R (5’-TTTCGCTGCGTTCTTCATCG-’3); and ITS86F (5’-GTGAATCATCGAATCTTTGAA-’3) and ITS4R (5’-TCCTCCGCTTATTGATATGC-’3), respectively, and the enzyme AmpliTaq Gold DNA Polymerase (Applied Biosystems, Foster City, CA, USA). The thermal cycling conditions were 95°C for 10 min then 95°C for 30 s, 55°C for 30 s, and 72°C for 1 min for 35 cycles. The amplicons obtained from each fecal sample were pooled and sequenced on IT PGM equipment (Life Technologies, Waltham, MA, USA) using a 318 chip.

#### Illumina sequencing

Samples were quantified using a fluorometer (Qubit), and 12.5 ng of DNA were used to amplify the bacterial 16S rRNA gene, specifically targeting the V3–V4 hypervariable regions. Amplification was performed using KAPA HiFi DNA polymerase (Roche, ref: 07958935001) and the following primer pair: 16S Amplicon PCR Forward Primer 5’ TCGTCGGCAGCGTCAGATGTGTATAAGAGACAGCCTACGGGNGGCWGCAG and Reverse Primer 5’ GTCTCGTGGGCTCGGAGATGTGTATAAGAGACAGGACTACHVGGGTATCTAATCC. These primers include overhang adapters compatible with Illumina sequencing. Library preparation was carried out using the Nextera XT DNA Library Preparation Kit (Illumina, ref: FC-131-1096, San Diego, CA, USA), which attaches dual indices and sequencing adapters. Sequencing was performed on an Illumina MiSeq platform using the MiSeq Reagent Kit v3 (2 × 300 bp), which provides sufficient read length to fully cover the amplified V3–V4 region of the 16S rRNA gene.

### Computational resources

For the creation of *Mbiome* and posterior running and validation analyses a single computing node with 62 GB of RAM and 6 cores was used for all analyses.

## Data availability

All the data generated and used for the comparative analysis is available at 10.5281/zenodo.17512479.

## Code availability

All the code for the comparative analysis is available at 10.5281/zenodo.17512322 and https://github.com/ZabalaAitor/ComparativeAnalysisOfMbiome. The *Mbiome* workflow is available at https://github.com/MGorostidi/mbiome.

## Author contributions

Author contributions are detailed according to the CRediT taxonomy. MGA, LM and DO contributed to the conceptualization of the study, as well as the designed of the methodology. Software development and implementation was carried out by MGA. Validation of results was conducted by MGA, AOC, AZ and LM. Formal analysis was performed by MGA and AZ. MGA, AOC and LM were responsible for the investigation. Resources were provided by LM and DO. Data curation was handled by MGA, AOC and LM. Original draft was mainly written by MGA, but all authors—MGA, AOC, AZ, LM, and DO—contributed to reviewing and editing the manuscript. Visualization was carried out by MGA and AZ. Supervision was provided by LM and DO, while project administration was managed by MGA, LM and DO. Funding acquisition was undertaken by LM and DO.

## Acknowledgments

The authors have used generative AI technology (ChatGPT) to improve the readability and overall quality of the manuscript. The use of AI has not altered the content or messages conveyed in the manuscript, focusing solely on refining the writing for clarity and enhanced readability.

